# Climate and competition combine to set elevational distributions of tropical rainforest *Drosophila*

**DOI:** 10.1101/2022.04.01.486700

**Authors:** Jinlin Chen, Owen T. Lewis

## Abstract

Species turnover with elevation is a widespread phenomenon and provides valuable information on why and how ecological communities might reorganize as the climate warms. Tropical mountains typically have pronounced thermal gradients and intense species interactions, providing a testing ground for investigating the relationship between thermal tolerances and biotic interactions as the proximate factors influencing species’ distributions. We investigated temperature and interspecific competition as causes of species turnover and abundance changes of the nine most abundant species of *Drosophila* along elevational gradients in the Australian Wet Tropics. Thermal performance curves revealed that species’ distributions were better explained by their performance at extreme temperatures, rather than their thermal optima. Upper thermal limits varied less among species than lower thermal limits. Nonetheless, these small differences were associated with differences in centred elevation of distribution, consistent with environmental sorting as a driver of community composition at low-elevation sites. In contrast, community composition at cool, high elevations was driven by temperature-dependent interspecific competition rather than tolerance to low temperatures. These results run counter to common assumptions about the role of abiotic and biotic factors in structuring communities along thermal gradients, and indicate that tropical insects may be highly vulnerable to future warming. Our study illustrates the importance of experimental, quantitative tests across biological levels (i.e., individuals to populations) and temporal scales (i.e., within-generation to multi-generation) for characterizing effects of climate on a guild of closely-interacting species.

## INTRODUCTION

Temperature has a fundamental impact on the reproduction, survival, growth, and behaviour of organisms (Huey and Kingsolver 1989; Huey and Stevenson 1979), strongly influencing species’ ranges and abundances (Hoffmann and Blows 1994; Wilson et al. 2005). As a result, estimates of thermal tolerances based on laboratory assays or species’ distributions have been used widely to evaluate species’ sensitivity to climate change (Deutsch et al. 2008; Kearney and Porter 2009). Tropical ecosystems comprise many species that live close to their upper thermal limits (Deutsch et al. 2008; Diamond et al. 2012; Huey et al. 2009), and which may not tolerate or adapt to warmer temperatures (Bonebrake and Deutsch 2012; Kellermann et al. 2012). The narrow thermal ranges of tropical insects (Khaliq et al. 2014) also mean that they will need to undertake relatively large latitudinal or elevational range shifts to track their climate envelopes as the climate warms, increasing the risk of extinction and community disassembly (Colwell et al. 2008; Sheldon, Yang, and Tewksbury 2011).

Despite high relevance for understanding species’ responses to climate change in tropical ecosystems (Corlett 2012), it remains questioned whether temperature is the major proximate factor setting the position of the ‘warm’ (low-latitude or low-elevation) limits of species’ distributions. Global analyses found smaller distribution shifts in response to warming at warm range limits than at cool range limits (Chen et al. 2011; Sunday, Bates, and Dulvy 2012), and some laboratory assays have found similar upper thermal limits for tropical and temperate species (MacLean et al. 2019; Nowrouzi et al. 2018; Overgaard, Kearney, and Hoffmann 2014). Instead, other abiotic factors (e.g. precipitation) or biotic interactions may be critical in setting range limits in the tropics (Engelbrecht et al. 2007; Louthan, Doak, and Angert 2015).

A common expectation, first expressed by Charles Darwin, is that tolerance to low temperatures will set cool range limits, while biotic interactions will more often set warm range limits (O’Brien et al. 2017; Paquette and Hargreaves 2021). However, a recent analysis of data for 654 taxa found that biotic factors became less important in determining warm limits towards the tropics, while abiotic factors remained consistently important across latitudes (Paquette and Hargreaves 2021), highlighting the regional differences in the relative roles of abiotic and biotic factors on local thermal gradients.

There remains limited empirical evidence on the proximate factors setting range limits in species-rich tropical communities (Feeley, Stroud, and Perez 2017; Jankowski et al. 2013), and most such studies examine only the correlation between thermal traits (such as critical temperatures and optimal temperatures) and species’ distributions (Amundrud and Srivastava 2020; Cahill et al. 2014; García-Robledo et al. 2016; von May et al. 2017; Nowrouzi et al. 2018; Pintanel et al. 2021). A significant correlation between tolerance and distribution does not rule out a role for biotic interactions: differing sensitivities of reproductive performance to temperature can further increase the competitive advantage of tolerant species over less tolerant species, sometimes driving qualitative changes in persistence (Lyu and Alexander 2022). Additionally, the abundance and distribution of a species may be sensitive to temperature change because of the thermal sensitivities of competitors, hosts, consumers, and other interacting species (Gifford and Kozak 2012; Merrill et al. 2008; Van Der Putten, Macel, and Visser 2010). In studies of species’ distributions along thermal gradients, niche underfilling has been often used to indicate the influence of biotic interactions (Chick et al. 2020; O’Brien et al. 2017), but biotic interactions such as competition, have rarely been measured directly. A coupled characterization of fundamental thermal performance and temperature-dependent biotic interactions within a complete guild along an environmental gradient is lacking. Such studies can help unify the long-separate concepts of environmental and biotic filters (HilleRisLambers et al. 2012), as well as provide realistic information on thermal sensitivities to inform conservation efforts in the face of climate change.

To investigate the roles of thermal tolerances and biotic interactions in influencing species’ range limits and structuring communities, we focused on the community of *Drosophila* flies occupying mountain rainforest habitats in northeastern Australia. These and other tropical mountains provide natural environmental gradients to test the sensitivity of species to temperature (Corlett 2011), with pronounced changes in species composition with elevation for many taxa (Williams, Bolitho, and Fox 2003). We hypothesized that species turnover along the elevational gradient would result from thermal constraints at cool, high-elevation sites, but from competitive exclusion at warmer, lowland sites. We first quantified species’ elevational distributions in the field, and then measured thermal performances and competitive outcomes to test the following predictions. (1) Species with relatively low abundance at upland sites (“lowland species”) would be the least cold-tolerant, but (2) species with relatively low abundance at lowland sites (“upland species”) would not be the least heat-tolerant. We further predicted that (3) species are competitively excluded from local assemblages by locally-abundant species under the thermal conditions typical at low elevations, but not under high elevation conditions.

## METHODS

We collected data on species’ elevational distributions in the field; their reproductive performance on thermal gradients and adults’ resistance to acute thermal stress (by knockdown assay); and the short-term and long-term outcomes of interspecific competition under low and high temperatures. All statistics mentioned in this section were performed with R version 4.0.3 (R Core Team, 2020). Original data are archived on Zenodo (10.5281/zenodo.6400611) and code for analyses is available on GitHub (https://github.com/Jinlinc/DistributionRegulators.git).

### Species distributions along elevational gradients

We used a standardized method to survey the relative abundance of different *Drosophila* species, allowing us to calculate indices of elevational distributions. These were used in later sections to test whether thermal traits correlate with distribution patterns.

#### Field survey

Field data on *Drosophila* communities were collected from rainforest sites at Paluma Range (S18° 59.031’ E146° 14.096’) and Kirrama Range (S18° 12.134’ E145° 53.102’), Queensland, Australia. These sites lie within the Wet Tropics bioregion which has high levels of endemism associated with cool and moist upland refugia (Williams et al. 2003). Abundances of *Drosophila* at these sites peak from March to June. *Drosophila* pupae were sampled using bottle traps baited with fermented banana from 11^th^ March – 12^th^ April 2016 for three sites at elevations of 70m, 350/390m, and 730/880m (subsequently referred to as low-, mid- and high-elevation, respectively) on each of the two mountains. At each site, we randomly sorted 182 *Drosophila* pupae from 15 traps (five traps of 12, 15 or 24-day exposure time) hanging at the understory level. From the 1092 pupae sent for sequencing, 716 pupae in total were successfully identified to species by DNA metabarcoding, with 86 – 134 pupae at each site (Jeffs et al. 2021). Two infrequent species, *Drosophila serrata* (1 pupa) and *D. immigrans* (4 pupae), were excluded from analyses.

#### Distribution analysis

Two methods were used to assess elevational distribution for individual species. First, the probability of detecting a particular *Drosophila* species among other co-occurring *Drosophila* species was modelled as a function of *elevation*, *mountain* and their interaction in a generalized linear model (family = “binomial”). For each species, the response variable was 1 if the pupa was identified as the focal species and 0 if it was identified as any other species. Second, since none of the species had a bimodal distribution with elevation, abundance-weighted mean elevation (*hIndex*) was used as a simple alternative method to quantify distributions. The location of each sample was assigned a value of 0, 0.5, or 1 if it was collected at low-, mid- or high-elevation sites, respectively. *hIndex* was calculated for each species by averaging these values for samples from both mountains. Correspondence between the two measures was tested by Spearman’s rank correlation test between the coefficient value for *elevation* in the regression and *hIndex*.

### Maintenance of laboratory cultures

To minimize adaptation to laboratory conditions, we maintained the *Drosophila* species by isofemale lines and then crossed them into mass-bred lines (MBLs) shortly (4-6 generations) before measuring thermal traits. The isofemale cultures were established in 2017 and 2018 from adults collected from high- and low-elevation sites. Cultures were maintained initially at 24°C at the Biology Centre, Czech Academy of Sciences, and then transferred to the Department of Zoology, University of Oxford, UK, in December 2018, where they were maintained at 25°C. Cultures and all experiments mentioned below were maintained under a 12h/12h light/dark cycle. There were approximately 15 to 30 non-overlapping generations in the Czech Republic and four to seven non-overlapping generations in Oxford before establishing mass-bred lines.

Each MBL is made of four isofemale lines (except for *D. pandora*, where only three isofemale lines were available). Previous studies have found no local adaptation to elevations and high gene flow between mountains (O’Brien et al. 2017; Schiffer et al. 2007). To reflect the average value of traits, the four isofemale lines were from different mountains and elevations if possible (Supplementary Table 1). A large population of each species was initiated using two independently-reared MBLs. These mass-bred populations were maintained at 25°C until 2020, and at 23°C thereafter. Measurements taken from these mass-bred populations should not have been influenced by maternal effects, acclimation, or isofemale line effects.

Eight Drosophila species (D. bipectinata, D. birchii, D. bunnanda, D. pallidifrons, D. pandora, D. pseudoananassae, D. simulans, D. sulfurigaster) sourced from the Queensland tropical rainforest, plus a laboratory population of D. melanogaster (wild type, Dahomey strain) were measured for cold resistance, heat resistance and thermal performance curves (collectively called thermal traits). Drosophila melanogaster does not occur naturally at the study sites, but was measured as a benchmark for future comparisons. Competitive outcomes under different temperature regimes were assessed for five species (D. bipectinata, D. pallidifrons, D. pandora, D. pseudotakahashii, D. sulfurigaster). Drosophila rubida was not measured because it was difficult to raise to a large number and to synchronize with the other species. The thermal traits of D. pseudotakahashii were not measured because its population was contaminated by another species at the time. A new mass-bred population crossed by two uncontaminated isofemale lines was constructed later for the competition experiment.

### Reproductive thermal performance

We exposed adult *Drosophila* to seven temperatures ranging from 14°C to 32°C and measured how their reproductive success changed with temperature (procedures are detailed in Supplementary Figure 1A). We extracted reproductive thermal traits from the thermal performance curves and examined their correlations with centred elevations of distribution.

#### Experimental measurements

To generate the adult flies for measurements, fly eggs collected from the population cage were reared at low density (<100 eggs per vial) at 25°C. Emerging adults were separated by sex within 12 hours to guarantee that they were unmated. Rearing was started on different days for different species to synchronize the first day of egg-laying. Two additional vials with five pairs of flies were monitored daily for sexual maturation, indicated by egg laying. Two days after the first observation of egg-laying in both vials, two virgin females and two virgin males were paired in a new vial containing 4ml *Drosophila* medium (weight/volume concentration: 8% corn flour, 4% yeast, 5% sugar, 1% agar, and 1.67% methyl-4-hydroxybenzoate). Vials were randomly assigned to water baths set at one of seven constant temperatures (14°C, 17°C, 20°C, 23°C, 26°C, 29°C, 32°C). This range of temperature spans the temperatures our *Drosophila* usually experience in the field. For each species and each temperature treatment, eight replicates were evenly split between two experimental blocks, making up a total of 1512 vials. To measure productivity at different temperatures, vials were submerged with the water level kept above the zone within which flies could freely move. The temperature and relative humidity of vials in each water bath was monitored within two empty tubes placed at the centre and the corner. Observed temperatures varied ±0.5°C around the set temperature. Temperatures in the centre of the water bath were on average 0.5°C higher than at the corners; the average of the former during the experimental period was used as the corrected temperature in analyses. Relative humidity levels were similar to field conditions, ranging between 80% - 95%.

Flies in the experimental vials were left to lay eggs on fresh food for 48 hours (vials from the 1^st^ and 2^nd^ days). They were then transferred to new food and kept in their corresponding water baths for a further four days before being put onto new food for another 48 hours (vials from the 7^th^ and 8^th^ days). Vials containing eggs from the two 48-hour periods were kept at the corresponding temperature until adults emerged. Productivities for the two periods are shown separately in Supplementary Figure 1B. We averaged these values to reflect the relative reproductive success in early adult life (within 8 days of sexual maturity).

After eight days in water baths, all flies were returned to 25°C, a favourable temperature, in new vials with fresh food for a further four days before being sacrificed. Their offspring developed at 25°C until eclosion. These adult offspring numbers reflect the recovered reproduction following eight-day exposure to different temperatures. The numbers of surviving parents were recorded when flies were transferred onto new food on the 3^rd^, 7^th^, 9^th^, and 13^th^ day. Offspring numbers were counted 5 – 7 days after the first emergence was observed. The experiment was conducted from May to August 2019.

#### Thermal performance curves

A multi-level, non-linear piecewise model was fitted to describe how reproductive success changed with temperature for all *Drosophila* species. We used the Briere2 function (Briere et al. 1999) to describe how reproductive performance (square-rooted daily productivity per female) changed with temperature:

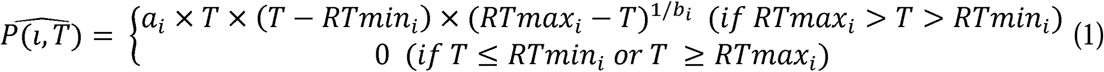

where 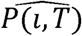 is the theoretical reproductive performance of species *i* at temperature *T*, *RTmin_i_* and *RTmax_i_* are the minimum and maximum temperatures for species *i* to reproduce, *a_i_* is a scaling factor, and *b_i_* is a shape factor of the curve for species *i*.

The observed reproductive performance, *P*(*i,T*), was modelled assuming a Gaussian distribution of errors as shown in Equation (2). A Gaussian distribution is not ideal to model transformed count data, which are all positive. However, models using untransformed counts with Poisson, zero-inflated Poisson, negative binomial, or lognormal distributions did not adequately converge.

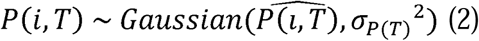

As standard deviations, *σ_σ_P__*_(*T*)_^2^, tend to get smaller as temperature approaches the critical points (beyond which performance is zero), we assumed seven different *σ_P_* _(*T*)_ for our seven test temperatures. They follow:

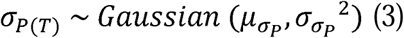

where mean, *µ_σ_P__*, and variance, *σ_σ_P__*^2^, are hyperparameters that describe the mean and variance of *σ_P_* _(*T*)_.

*RTmin_i_*, *RTmax_i_*, *a_i_* and *b_i_* of the Briere2 function are features of *Drosophila* species. We assumed that each value showed some similarities as they are all *Drosophila* species. To reflect this structure, *RTmin_i_*, *RTmax_i_*, *a_i_* and *b_i_* are modelled by equations (4) – (7). Their means and variances are also hyperparameters defined by priors. Additionally, the values of *a_i_*and *b_i_* were bounded to be positive. The values of *RTmin_i_* were bounded to be lower than 17°C and the values of *RTmax_i_* were bounded between 26°C to 35°C based on prior knowledge of the range of temperatures under which our species can reproduce.

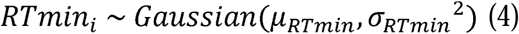

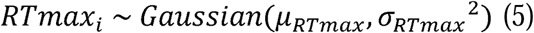

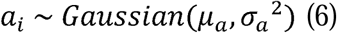

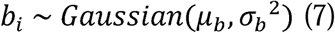

Prior distributions of hyperparameters *µ_RTmin_*, *µ_RTmax_*, *µ_a_*, *µ_b_*, and *µ_σ_P__* are Gaussian distributions with reasonable means (15**°**C, 30**°**C, 0, 0, 0) and large standard deviations (10, 10, 1, 10, 10) compared with the realistic range of these parameters in our species. The prior distribution of the hyper parameters *σ_RTmin_*^2^, *σ_RTmax_*^2^, *σ_a_*^2^, *σ_b_*^2^, and *σ_σ__p_*^2^ is inverse-gamma (0.001, 0.001), which is a commonly used non-informative distribution for prior of variance.

The multi-level model was fitted under a Bayesian framework using MCMC sampling by the *rstan* package (Stan Development Team 2021). Models converged and performance was acceptable in diagnostic plots (Supplementary Figure 2). Medians of the posterior distributions for *RTmin_i_*, *RTmax_i_*, *a_i_*and *b_i_* were used to construct the thermal performance curve of species *i*. Optimal temperatures for peak reproduction (*RTopt*) were deduced from known parameters:

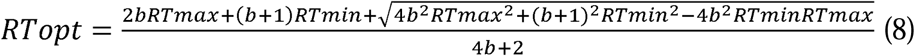

#### Testing for associations between reproductive thermal traits and species distributions

Among the tropical *Drosophila* species, the medians of the posterior distributions of *RTmin*, *RTmax*, and *RTopt* were modelled as a function of *hIndex* in a linear model with phylogenetic correction. The phylogenetic tree was derived with modification from Finet et al. 2021 (Supplementary Figure 3). Productivity at 29°C and 17°C, and recovered productivity after 29°C and 14°C, were used as direct measurements of performance at high or low temperatures. These temperatures were chosen because they are known to cause stress to the reproduction of the focal *Drosophila* species, and correspond to realistic understory air temperatures during the hottest and coolest months. These productivities were modelled as a function of *hIndex* and *experimental block* as fixed effects, and *species* as a random effect in generalized linear mixed-effect models (family = “negative binomial”) with phylogenetic correction, using the *brms* package (Bürkner 2021). *Drosophila simulans* has not been found in the standardized assay, therefore, it was not included in the regression. It is potentially a lowland-biased species because we only found their isofemale lines from lowland sites when we tried to collect as many isofemale lines as possible from different elevations. Adding *D. simulans* as a lowland-only species into the regression will not change the conclusions.

### Thermal knockdown

We measured the physiological tolerance of each *Drosophila* species to extremely cold and hot environments, and then examine their correlations with distribution patterns.

#### Experimental measurements

Resistance to extreme cold temperature was measured as knockdown time for each individual at 5°C and the time for recovering muscle coordination after a 30-minute exposure to 5°C. The constant temperature chosen for cold stress studies is often around 0°C (Gibert et al. 2001). As tropical species often have significantly weaker cold resistance (Gibert et al. 2001), 5°C was used instead to increase the variation among the tested species after pilot trials. Heat stress was chosen to be 40°C, which follows common practice for *Drosophila* species (Hoffmann, Sørensen, and Loeschcke 2003) and is expected to capture the between-species variance in heat tolerance over a timescale which is convenient to measure (Jørgensen, Malte, and Overgaard 2019). After being knocked down by heat (40°C), most flies did not survive. In this case, only knockdown time was used to evaluate resistance to heat. The knockdown experiments were conducted from May to Jun 2019.

Virgin adult flies (siblings of those used for reproduction measurements) were kept in same-sex groups at 25°C for 9-10 days before knockdown assays. Assays were conducted for male and female flies separately. An observation rack was divided into nine (3×3) cells. Each cell was randomly assigned one of the nine *Drosophila* species and held seven flat-bottomed 3ml glass vials, each with a randomly-selected individual of the allocated species. One set of observations on such a set-up represents a single block. We repeated measurements for three experimental blocks, and the allocation of species to cells was redrawn for each block. In total, we measured 21 individuals per sex per species. During measurement, the observation rack was moved immediately into the incubator pre-set at 5°C or 40°C. Every tube was examined once every minute and flies that lost their ability to stand in that minute were recorded. After exposure to 5°C for 30 minutes, all flies were in a chill coma. The observation rack was moved to a 25°C room. Flies were left undisturbed and the time taken until each fly regained its ability to stand was recorded.

#### Testing for associations between knockdown resistance and species distributions

The knockdown time by the heat, knockdown time by the cold and recovery time from the cold of each sex were first compared among species using ANOVA. When interspecific variation was observed, they were modelled as a function of *hIndex*, *block*, and *cell position* as fixed effects, and *species* as a random effect in linear mixed-effect models with phylogenetic correction using the *brms* package. *Species* was included as a random effect to account for repeated measures.

### Short-term competition

Pairs of *Drosophila* species were reared together in the same vials for one generation to evaluate how interspecific competition affects the observed reproductive success and predicted coexistence patterns under temperature regimes typical of low-elevation and high-elevation sites.

### Experimental design

We used incubators set at alternating temperatures mimicking average day-time and night-time temperatures in February at high-elevation (23°C / 21°C) and low-elevation sites (28.5°C / 24°C) (Supplementary Figure 4). Adults to establish the competition experiment were reared in bottles (30ml fresh medium) at moderate density (300 – 500 per bottle) at their testing temperatures. After eclosion, individuals that emerged within 48 hours were kept together in mixed-sex containers. Two days after the first observation of egg-laying, adults of different sexes were separated, and from the following day were allowed to pair and lay eggs in new experimental vials (5ml medium) for two days, without or with another competitor species.

Five species were chosen as representative species for upland-biased, elevation-generalist and lowland-biased distribution types (see Results for the definition of distribution types). The two-species combinations were lowland species vs. upland species: *D.bipectinata* vs. *D. pallidifrons*, *D.bipectinata* vs. *D. pseudotakahashii*, *D.pandora* vs. *D. pallidifrons*; lowland species vs. lowland species: *D. bipectinata* vs. *D.pandora*; lowland species vs. elevation-generalist: *D. bipectinata* vs. *D. sulfurigaster*; upland species vs. elevation-generalist: *D. pallidifrons* vs. *D. sulfurigaster*. Each combination was tested at different founding densities in a factorial design: (4 pairs of species A, 2 pairs of species B), (4A, 4B), (4A, 8B), (2A, 4B) and (8A, 4B). We also included monocultures of each species with 2, 4, and 8 pairs. Each density and species combination was replicated ten times across two or three blocks staggered by two days (two blocks for the *D. pandora* vs. *D. pallidifrons* combination; three blocks for the other five pairs). Offspring that successfully developed to adulthood were identified to species and counted. The experiment of *D. pandora* vs. *D. pallidifrons* was conducted from September to December 2020 and the other five pairs were experimented with from January to March 2021.

#### Short-term competitive outcomes

We used the Beverton-Holt model to describe the population growth of a single generation of flies on discrete and temporary resources:

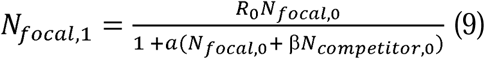

Where *N_focal_*_,0_ is the number of parents and *N_focal_*_,1_ is the number of offspring of the focal species. *N_competitor_*_,0_ is the number of parents of the competing species. *R_0_* is the generational reproduction rate and *a* is a constant defining the form of the density-dependence relationship. *Β* represents the interspecific competition coefficient of the competitor species to the focal species, and defines the equivalence between the two competing species. Offspring numbers of the focal species were modelled assuming a negative binomial error distribution, under a Bayesian framework using MCMC sampling by the *rstan* package, as described by Terry et al. (2021). Model diagnostics are shown in Supplementary Figure 5A-B. Raw data and fitted curves of each pair are shown in Supplementary Figure 6. Median values of posterior distributions were used as parameters to infer the equilibrium states of each pair following Hassell and Comins (1976).

### Long-term competition

To validate the long-term population-level outcomes of asymmetrical competition shown in the short-term competition experiment, we raised one lowland species and one upland species in monoculture and mixed-species culture for multiple overlapping generations under temperature regimes typical of low-elevation and high-elevation sites. We then evaluated the long-term effects of temperature and competition on population sizes.

#### Experimental design

Four monocultures of each species and eight mixed-species cultures were maintained at each temperature regime for 13 weeks, totalling 32 cultures. The cultures were evenly divided into two blocks starting on different dates. Monocultures were started with ten pairs of individuals. Mixed-species cultures were started with ten pairs of individuals of each species. The starting density was very low compared to the equilibrium density. Each culture was maintained in a series of five bottles (30ml fly medium) following Ayala, Gilpin, and Ehrenfeld (1973). At the start of each week, adults surviving in the most recent bottle and adults which were freshly emerged in the older four bottles were separately collected, photographed and transferred together into a new bottle with fresh food. In this way, adult survival and reproduction were recorded separately. As the total population sizes were relatively stable, they were only counted when the populations were terminated after 13 weeks of maintenance. To avoid pseudo-replication introduced by ‘incubator’ effects, the two incubators were switched between temperature regimes every week, with their contents moved accordingly. Trays were shuffled inside the incubator every two days. Temperature and humidity were recorded and the temperature regimes were confirmed during and at the end of experiments. The experiment was conducted from September to December 2020.

#### Testing temperature and competition effects on long-term population persistence

Population sizes were modelled as a function of *temperature*, *species*, *presence/absence of competitors*, and their interactions, with *culture ID* as a random effect in a generalized linear mixed-effect model (family = “zero-inflated negative binomial”) using the *brms* package. Model diagnostic is shown in Supplementary Figure 5C.

## RESULTS

### Field distributions

The numbers of samples found at low-, medium- or high-elevation sites for each of the nine major *Drosophila* species (accounting for 99% of all samples) are shown in Figure 1. Distributions quantified using regression and by centred elevation (*hIndex*) were consistent (Spearman’s rank correlation rho = 0.93, p value = 0.0007). Among the three elevations sampled, the detection probability of *D. bipectinata* and *D. pandora* increased monotonically towards lower-elevation sites and were thus categorized as lowland-biased species with high confidence. *Drosophila pseudoananassae* showed a lowland bias on one of the two mountains, and a mid-elevation peak on the other. *Drosophila rubida*, *D. sulfurigaster* and *D. birchii* showed no significant change with elevation and were thus defined as elevation generalists. *Drosophila pallidifrons* and *D. pseudotakahashii* were significantly more likely to be found at high elevations and were thus defined as upland-biased species. Coefficients and p values of the regressions and *hIndex* values are shown in Supplementary Table 2. For D*. bunnanda,* the six records were insufficient for model fitting but all occurred at lowland sites, and data from a large-scale study suggest that it is a lowland-biased species (Schiffer and McEvey 2006).

**Figure 1.**
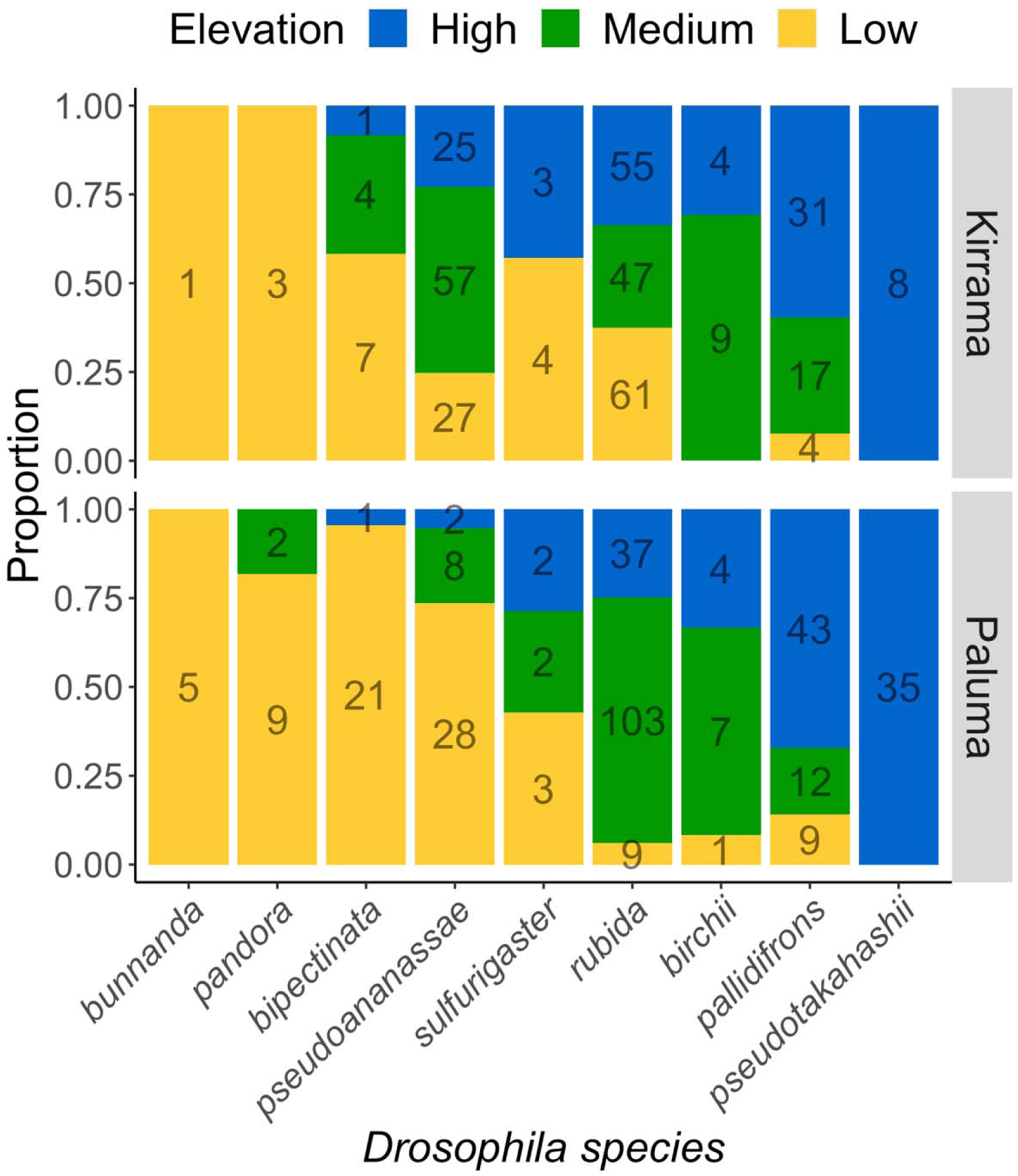
Distribution patterns of nine rainforest *Drosophila* species in Kirrama Range or Paluma Range. The proportion of samples found at sites at low (yellow), medium (green) and high (blue) elevations is shown for each species. Numbers on the bars show the counts of samples for each combination of species and site.

### Thermal performance curves

Thermal performance curves of daily productivity per female parent vary among species in terms of the range, optimal temperature, peak productivity, and shape factors (Figure 2; Table 1; see Supplementary Figure 2C for original data and fitted curves for each species). The temperature for optimal reproductive performance, *RTopt*, did not correlate with *hIndex* (Coefficient = 0.09, 95% CI (credible interval) = -2.83 – 3.01; Figure 2b). Cold tolerances, *RTmin*s, varied more among species than heat tolerances, *RTmax*s (standard deviation: 2.82 versus 1.01, respectively). There was no general trade-off between *RTmin* and *RTmax* (Spearman’s rank correlation test: p value = 0.10; Figure 2c). For example, *D. sulfurigaster* outperforms its upland-biased relative, *D. pallidifrons*, across the temperature range.

**Figure 2.**
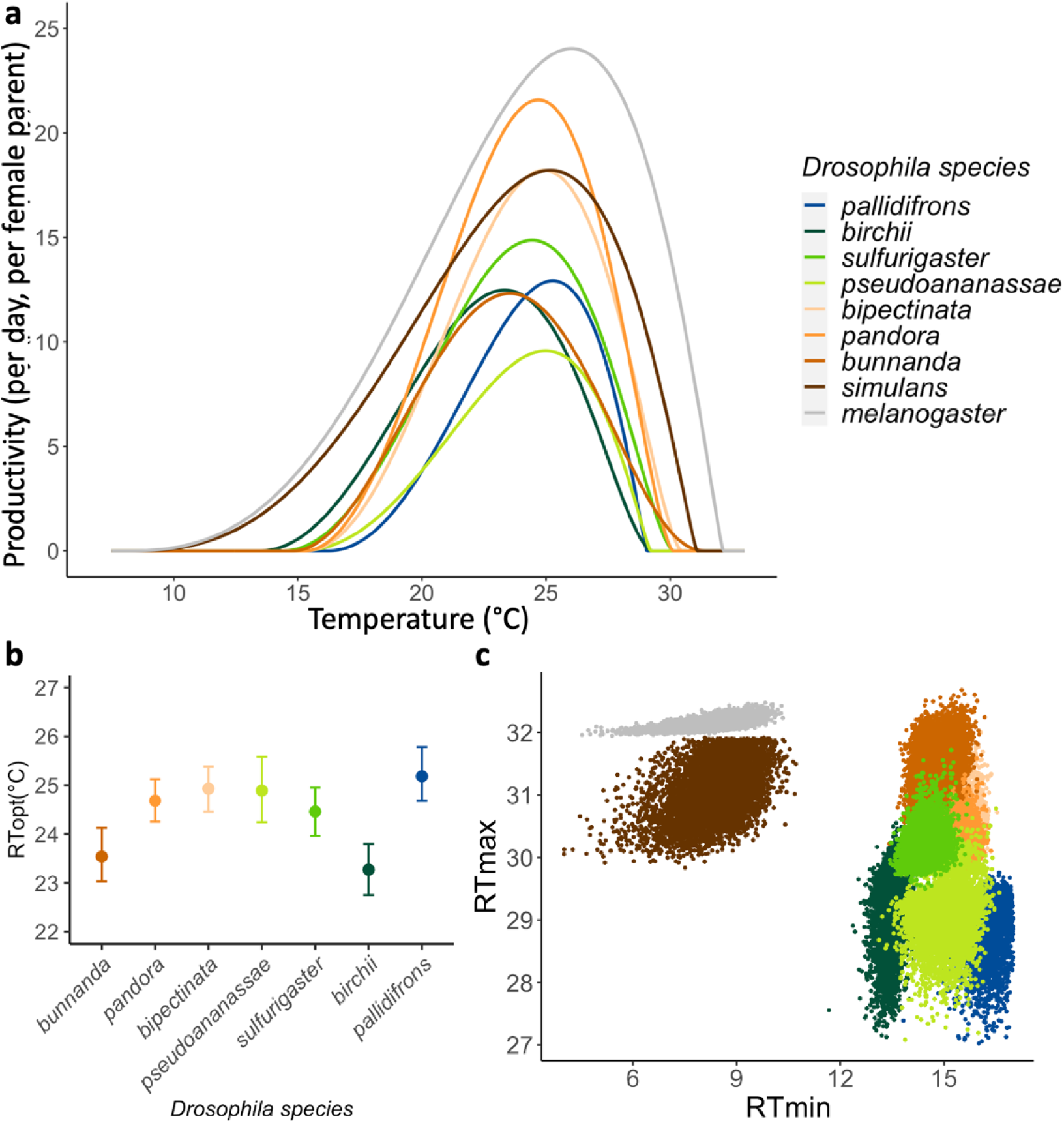
Reproductive thermal traits. (a) Thermal performance curves for reproduction of nine *Drosophila* species. The fitted adult offspring number (per female parent per day) are plotted against temperature. (b) Optimal temperatures for reproduction of the seven tropical *Drosophila* species, ordered by their *hIndex*. (c) Posterior distribution of *RTmax*s and *RTmin*s of each *Drosophila* species. Colours indicate distribution types: cold colours indicate upland-biased species represented and warm colours indicate lowland-biased species; the grey colour represents *D. melanogaster*.

**Table 1.** Parameters of thermal performance functions and their 90% credible intervals (ci90) of the nine species. RTmin and RTmax are the minimum and maximum temperatures for the species to reproduce. a is the scaling factor and b is the shape factor of the Briere2 function.

| species | a | a.ci90 | b | b.ci90 | RTmin | RTmin.ci90 | RTmax | RTmax.ci90 |
| --- | --- | --- | --- | --- | --- | --- | --- | --- |
| *D. bipectinata* | 0.0046 | 0.0031 - 0.0060 | 1.27 | 1.02 - 1.55 | 15.28 | 14.54 - 15.87 | 30.44 | 30.08 - 31.05 |
| *D. birchii* | 0.0034 | 0.0022 - 0.0056 | 1.17 | 0.95 - 1.56 | 13.45 | 13.07 - 13.77 | 29.27 | 28.12 - 29.81 |
| *D. bunnanda* | 0.0016 | 0.0012 - 0.0025 | 0.88 | 0.81 - 1.06 | 14.57 | 14.10 - 15.18 | 31.21 | 30.60 - 31.78 |
| *D. melanogaster* | 0.0037 | 0.0032 - 0.0042 | 1.72 | 1.48 - 2.03 | 8.29 | 6.89 - 9.37 | 32.13 | 32.03 - 32.28 |
| *D. pallidifrons* | 0.0073 | 0.0056 - 0.0098 | 1.74 | 1.37 - 2.38 | 16.23 | 15.54 - 16.75 | 29.06 | 28.19 - 29.38 |
| *D. pandora* | 0.0052 | 0.0037 - 0.0065 | 1.26 | 1.03 - 1.51 | 15.25 | 14.58 - 15.80 | 30.13 | 29.87 - 30.58 |
| *D. pseudoananassae* | 0.0053 | 0.0035 - 0.0071 | 1.68 | 1.24 - 2.32 | 15.06 | 14.17 - 15.89 | 29.22 | 28.38 - 29.77 |
| *D. simulans* | 0.0035 | 0.0027 - 0.0046 | 1.69 | 1.36 - 2.21 | 8.51 | 6.94 - 9.64 | 31.09 | 30.39 - 31.78 |
| *D. sulfurigaster* | 0.0040 | 0.0028 - 0.0051 | 1.27 | 1.03 - 1.52 | 14.37 | 13.91 - 14.96 | 30.11 | 29.84 - 30.63 |

### Testing correlations between cold tolerance and distribution

*RTmin*s were not correlated with species distribution patterns (Figure 3a. Coefficient = - 0.41, 95% CI = -4.15 – 3.43). Similarly, upland-biased species did not show higher productivity at the low temperature, 17°C (Figure 3b; Coefficient = -0.27, 95% CI = -3.80 – 3.05). When exposed to acute sublethal low temperature (5°C), all seven tropical *Drosophila* species showed similarly poor performance compared to *D. simulans* and *D. melanogaster* (Supplementary Table 3). Most species did not reproduce during their eight-day exposure to 14°C (Supplementary 1B), however, all of them were able to resume productivity once transferred back to 25°C. Species with a higher centred elevation of distribution had slightly but non-significantly higher recovered productivity (Figure 3c; Coefficient = 0.33, 95% CI = -0.55 – 1.14). It took longer, but not significantly so, for upland species to regain muscle coordination after chill coma (Figure 3d; Male: coefficient = 13.04, 95% CI = -9.31 – 35.05; Female: coefficient = 8.49, 95% CI = -2.18 – 20.02)

**Figure 3.**
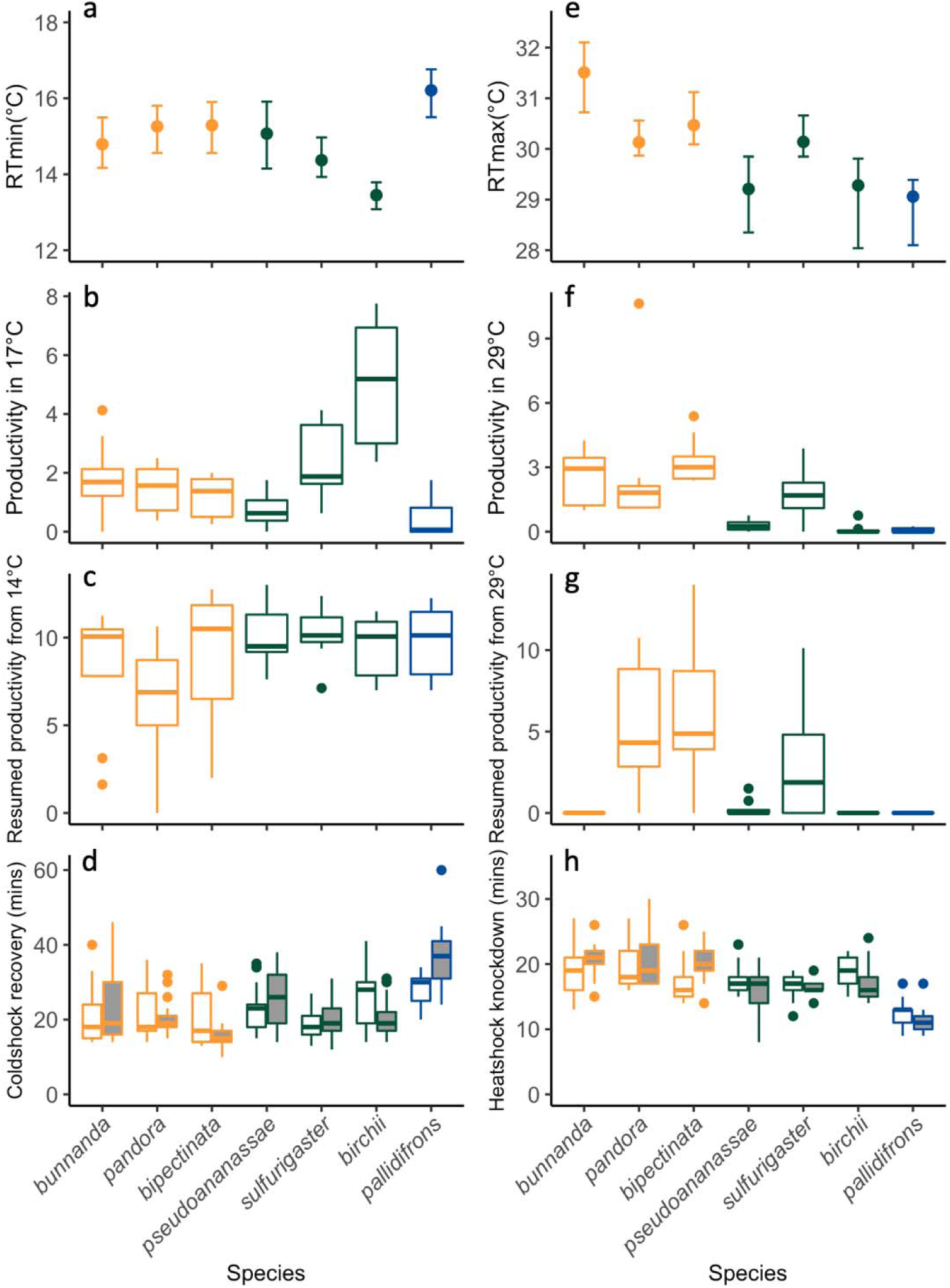
Reproductive and physiological thermal tolerance of seven tropical *Drosophila* species. Species are ordered by their *hIndex*, from lowland-biased species (left) to upland-biased species (right). Colours represent distribution types: orange for lowland species, green for elevation generalist and blue for upland species. Cold tolerance is represented by *RTmin* (a), productivity at 17°C (b), recovered productivity after 14°C (c) and recovery time after chill coma (d). Heat tolerance is represented by *RTmax* (e), productivity at 29°C (f), recovered productivity after 29°C (g) and knockdown time at high temperature (h). Graph (a) and (e) show the medians and 90% credible intervals of the posterior distribution of the estimated parameters. Boxplots in b-d and f-h show the minimum, 25^th^ percentile, median, 75^th^ percentile, maximum and potential outliers. In (d) and (h), data for females (shaded) and males (unshaded) are plotted separately.

### Testing correlations between heat tolerance and distribution

Species whose distribution were more biased towards lowland have higher critical temperatures for reproduction, *RTmax*s (Figure 3e; Coefficient = -3.06, 95% CI = -5.30 – -0.88). Reproductive performance at 29°C also decreased with *hIndex* (Figure 3f; Coefficient = -5.80, 95% CI = -9.37 – -2.50). After exposure to 29°C for eight days, the two species with the highest *hIndex* could not reproduce when transferred back to 25°C, while four out of the other five elevation-generalist and lowland-biased species resumed reproduction (Figure 3g). Knockdown time at lethally high temperature (40°C) was shorter among species of higher *hIndex* (Figure 3h; Male: coefficient = -6.71, 95% CI = -13.70 – 0.50; Female: coefficient = -1.99, 95% CI = -9.48 – 5.71), indicating these species lose their muscle coordination faster at high temperatures.

### Effects of interspecific competition at upland and lowland temperatures

When raised in a laboratory condition mimicking the warmer, lowland sites, reproductive success was highest for the two lowland-biased species, followed by the elevation-generalist species, *D. sulfurigaster*. The two upland species could barely reproduce regardless of the presence of competitors (Figure 4; Table 2: proliferation rates R_0_ < 1). *Drosophila pseudotakahashii*, whose distribution was most constrained to upland sites, had an even lower proliferation rate at high temperatures than *D. pallidifrons*. Their competitive effect on lowland species was minimal, indicated by low *β* values (Table 2). The two lowland species, *Drosophila pandora* and *D. bipectinata*, are expected to coexist stably in lowland, based on their reproductive and competitive parameters.

**Figure 4.**
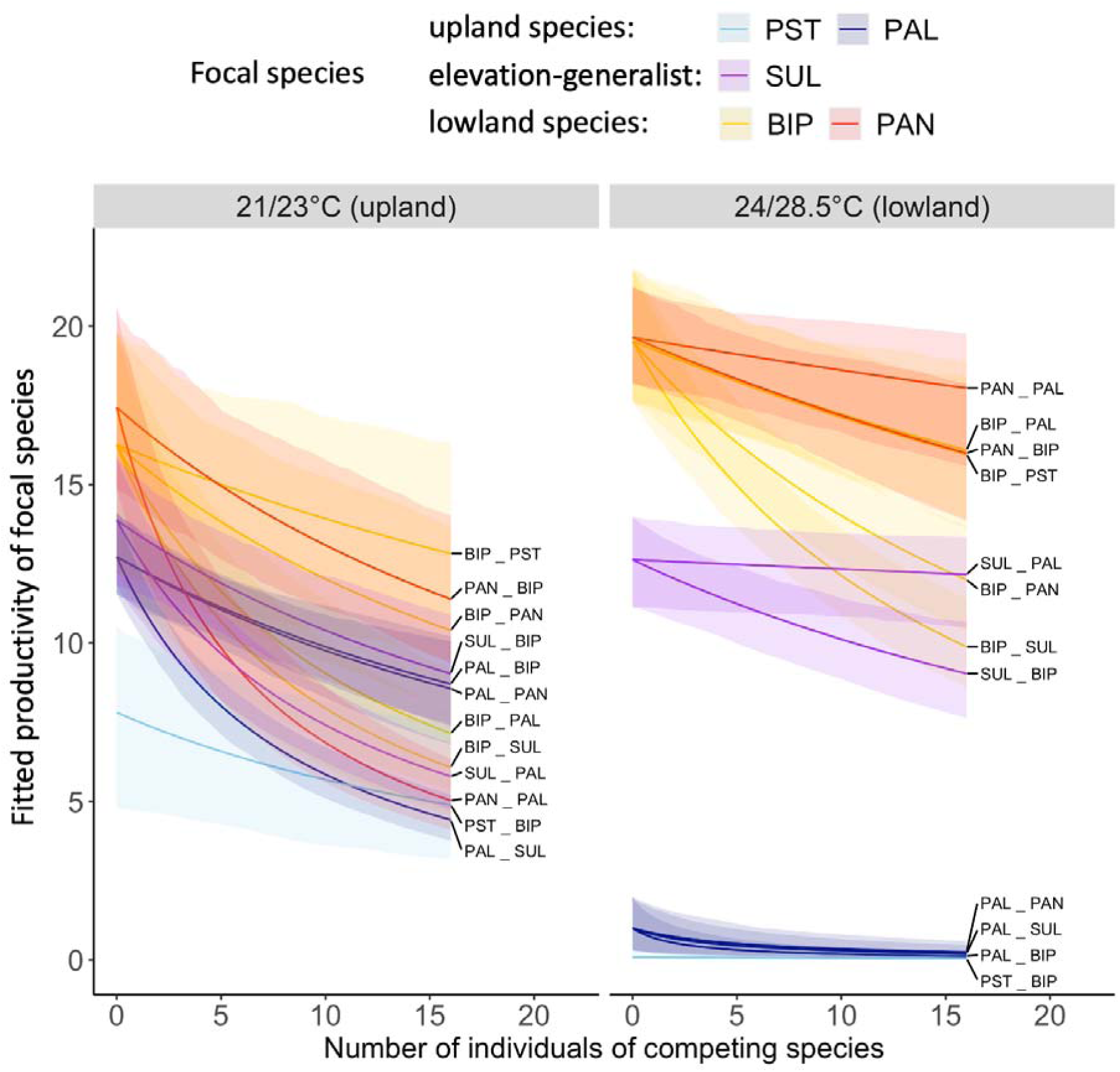
Interspecific competition for species pairs under upland (left) and lowland (right) temperature regimes. Each line shows the two-day productivity per pair of the focal species (four pairs in total) while changing the number of individuals of the competing species. The identity of the focal species in each pair is indicated by the colour of the line (see legend) and by the label (e.g., for BIP_PST, the first code BIP is the focal species and the competitor species is PST). The shaded area indicates the 90% credible interval of the fitted values under the Beverton-Holt model for pairwise species competition. PST = *D. pseudotakahashii*, PAL = *D. pallidifrons*, SUL = *D. sulfurigaster*, BIP = *D. bipectinata*, PAN = *D. pandora*.

**Table 2.** Fitted values of the parameters of the competition model and predicted equilibrium states for pairwise interspecific competition. R_0_ is the reproductive rate, a is a constant defining the form of the density-dependence relationship, f3 is the interspecific competition coefficient, and ci90 represents the 90% credible intervals of each parameter.

| 887 | Temperature | Focal species | $R_0$ | $R_0$.ci90 | $a$ | $a$.ci90 | Competitor | $\beta$ | $\beta$.ci90 | Equilibrium state of the focal species |
| --- | --- | --- | --- | --- | --- | --- | --- | --- | --- | --- |
|  | Cool, upland | BIP | 11.36 | 8.06-15.21 | 0.05 | 0.02-0.09 | PAL | 2.26 | 1.4-4.08 | excluded |
|  |  |  |  |  |  |  | PAN | 0.99 | 0.47-1.95 | excluded |
|  |  |  |  |  |  |  | PST | 0.47 | 0.12-1.09 | stable coexistence |
|  |  |  |  |  |  |  | SUL | 2.95 | 1.91-5.25 | excluded |
|  |  | PAL | 27.94 | 19.52-38.66 | 0.42 | 0.27-0.64 | BIP | 0.3 | 0.15-0.49 | dominant |
|  |  |  |  |  |  |  | PAN | 0.32 | 0.14-0.52 | dominant |
|  |  |  |  |  |  |  | SUL | 1.22 | 0.9-1.62 | unstable coexistence |
|  |  | PAN | 13.68 | 10.4-17.85 | 0.07 | 0.04-0.12 | BIP | 0.74 | 0.33-1.36 | dominant |
|  |  |  |  |  |  |  | PAL | 3.41 | 2.26-5.59 | excluded |
|  |  | PST | 6.27 | 3.4-10.66 | 0.08 | 0.03-0.19 | BIP | 0.79 | 0.35-1.76 | stable coexistence |
|  |  | SUL | 20.96 | 14.27-31.13 | 0.25 | 0.14-0.44 | BIP | 0.41 | 0.19-0.67 | dominant |
|  |  |  |  |  |  |  | PAL | 1.05 | 0.71-1.53 | unstable coexistence |
|  | Warm, lowland | BIP | 15.35 | 12.51-19.05 | 0.07 | 0.05-0.11 | PAL | 0.29 | 0.07-0.63 | dominant |
|  |  |  |  |  |  |  | PAN | 0.87 | 0.54-1.35 | stable coexistence |
|  |  |  |  |  |  |  | PST | 0.31 | 0.07-0.63 | dominant |
|  |  |  |  |  |  |  | SUL | 1.35 | 0.93-2 | excluded |
|  |  | PAL | 0.99 | 0.19-2.37 | 0.12 | 0.02-0.46 | BIP | 6.81 | 2.98-22.27 | unable to establish |
|  |  |  |  |  |  |  | PAN | 2.99 | 1.52-8.93 | unable to establish |
|  |  |  |  |  |  |  | SUL | 3.98 | 1.77-12.83 | unable to establish |
|  |  | PAN | 17.18 | 14.2-21.24 | 0.09 | 0.06-0.14 | BIP | 0.27 | 0.08-0.51 | stable coexistence |
|  |  |  |  |  |  |  | PAL | 0.11 | 0.01-0.29 | dominant |
|  |  | PST | 0 | \ | \ | \ | BIP | \ | \ | unable to establish |
|  |  | SUL | 13.65 | 10.26-19.03 | 0.15 | 0.09-0.24 | BIP | 0.37 | 0.18-0.62 | dominant |
|  |  |  |  |  |  |  | PAL | 0.04 | 0-0.14 | dominant |

When raised under a cooler, upland condition, all species could reproduce and sustain their populations (Table 2: all R_0_s were higher than 1). Lowland species were strongly affected by the density of *D. pallidifrons*, an upland species (shown by large *β* values), while upland species were significantly less affected by lowland species. Competition with *D. pallidirons* under upland conditions was predicted to drive *D. pandora* and *D. bipectinata* to exclusion (Table 2).

In the long-term competition experiment (Figure 5), high temperature drove the upland-species *D. pallidifrons* to extinction both in monocultures (except in one replicate, where two adults were found), and when reared together with the lowland competitors. In contrast, the monoculture of the lowland species *D. pandora* remained abundant at both low and high temperatures (temperature’s effect: 95% CI = -0.26 – 0.12, overlapping with zero). The presence of *D. pandora* only slightly decreased the population size of *D. pallidifrons* (median coefficient of competition = -0.32, 95% CI = -0.14 – -0.49). The presence of *D. pallidifrons* had no significant effect on the population size of *D. pandora* in warm, lowland temperatures (95% CI = -0.28 – 0.06), but had a strong negative effect (median coefficient of competition = -1.56, 95% CI = -1.73 – -1.40) in cool, upland temperatures.

**Figure 5.**
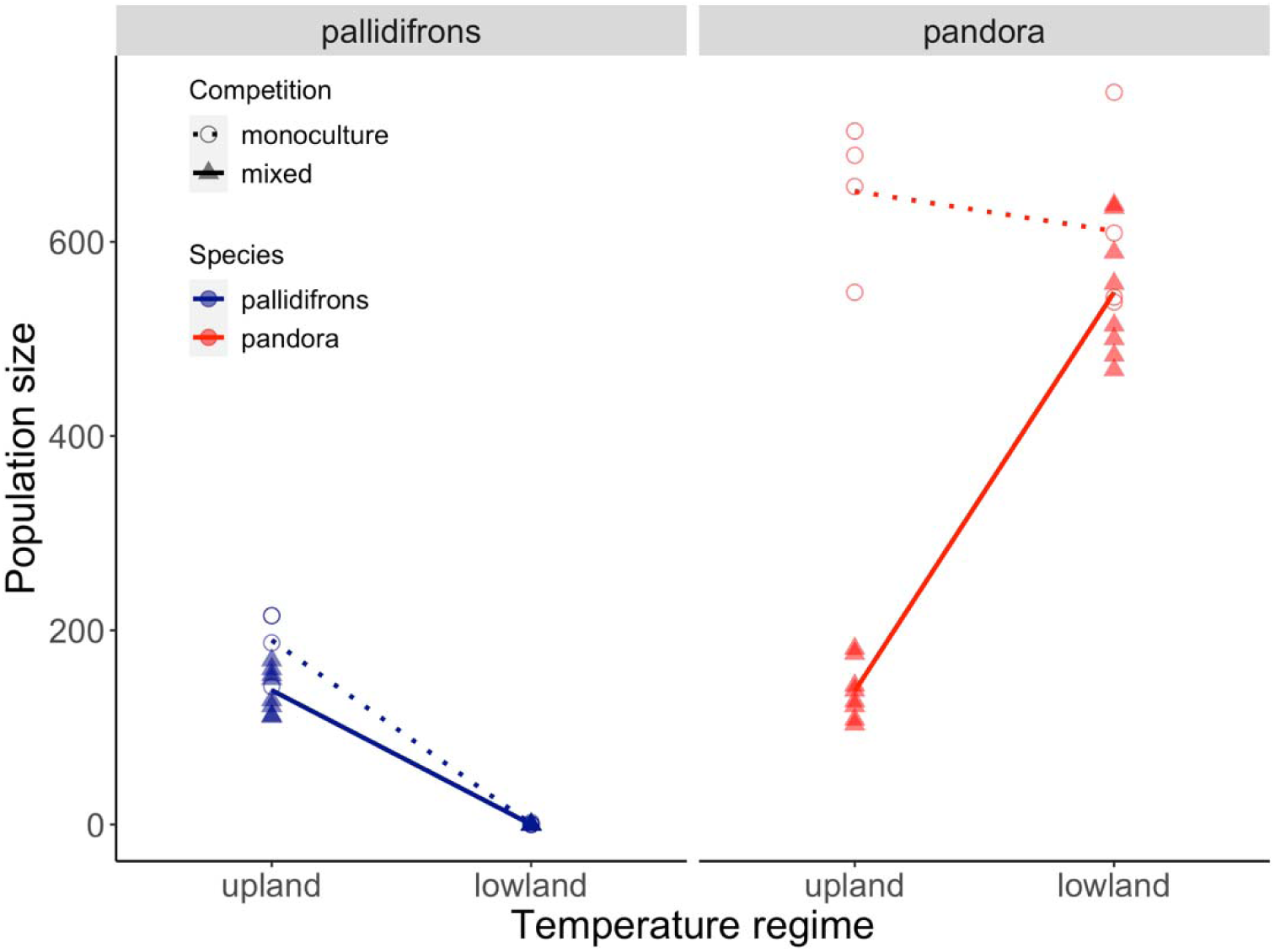
Effects of temperature and interspecific competition on population sizes of *D. pallidifrons* (upland species, blue) and *D. pandora* (lowland species, red). Each point indicates the ending population size of each culture (n(monoculture) = 4, n(mixed-species) = 8). Dashed (monoculture) and solid (mixed-species culture) lines connect the means of population sizes in the two temperature regimes.

## DISCUSSION

Our results do not support the common assumption that cool boundaries to species’ ranges are constrained abiotically (reflecting thermal niches), while biotic interactions (such as interspecific competition) define warm boundaries. Our study adds to a growing literature for tropical mountain systems showing significant associations between upper thermal limits and patterns of species turnover with elevation (Amundrud and Srivastava 2020; García-Robledo et al. 2016; von May et al. 2017; Pintanel et al. 2021). Noteworthy, this result does not rule out the potential contribution of competition with warm-adapted species to raising the warm boundaries to higher elevations. However, the observation that upland species were excluded from the lowest elevations solely by high temperatures experienced on a daily basis implies that further warming will likely drive contraction of their ranges directly. In contrast, lowland species were not worse than upland species at surviving or reproducing at low temperatures. However, lowland species were outcompeted by one upland species whose distribution was confined to high elevations as a result of intolerance to heat. This important role of interspecific competition at cold boundaries on elevational gradients (also shown by Gifford and Kozak 2012; Lyu and Alexander 2022; Rodrıguez-Castañeda et al. 2017) helps to explain the maintenance of species richness and endemism in mountain regions.

### Abiotic versus biotic limits of distribution

The idea that contributions of abiotic versus biotic factors might differ for species’ warm and cool range boundaries can be traced back to Charles Darwin, and remains an area of active debate (Cahill et al. 2014; Hargreaves, Samis, and Eckert 2014; Schemske et al. 2009). There is especially mixed evidence on the importance of upper thermal limits in setting warm boundaries (positive relationships: Batista et al. 2018; Duarte et al. 2012; García-Robledo et al. 2016; Kellermann et al. 2012; Merrill et al. 2008; null relationships: Brandt et al. 2020; Gaston and Chown 1999; Huang and Tu 2008; Kimura 2004; Nowrouzi et al. 2018).

This inconsistency could arise for at least three reasons. First, data available for synthetic studies over-represent systems from temperate latitudes in the northern hemisphere (Feeley et al. 2017; Parker 2022) and relate largely to cool limits (Cahill et al. 2014). Comparative studies have shown in lower latitudes that local variations in thermal tolerance have a tighter association with environmental variations (Duarte et al. 2012), and that species distribution appears more sensitive to warming (Freeman et al. 2021). Second, mechanisms governing latitudinal or elevational distribution patterns could differ. Studies of *CTmin* (the critical temperature at which organisms lose their muscle coordination) have found a tight relationship between this lower thermal limit and species’ latitudinal distributions (Kellermann et al. 2012; Overgaard et al. 2014). In contrast, we found that various cold tolerances of our species, including *CTmin* sourced from the literature, were not indicative of elevational distribution patterns. Third, some of the uncertainty might reflect methodological differences when choosing, defining and measuring thermal traits.

Different choices of indicators of thermal performance, and different experimental methods may explain why some of our results contrast with some other studies of Australian rainforest *Drosophila*. Overgaard et al. (2014) found no evidence that the upper thermal limits of tropical rainforest *Drosophila* differed among species. Unlike Overgaard et al. (2014), we measured parental productivity by allowing their eggs to develop into adults under the relevant temperature. We observed that many eggs laid at high temperatures never hatched, perhaps as a result of sperm sterilization (Parratt et al. 2021). We also observed different speeds of senescence after exposure to different temperatures (Supplementary Figure 1B), which the three-day period of fecundity measurement in Overgaard’s study would not have captured. Nevertheless, both studies highlight the conserved nature of heat tolerance and suggest small thermal safety margins for future warming.

Field experiments on *D. birchii* (O’Brien et al. 2017) found that its single-generation reproduction in monoculture was highest at low elevations. In our experiments, *D. birchii* was one of the most heat-sensitive species, and its populations were unable to persist within a few generations at high temperatures typical of low elevations (pilot experiment, unpublished data). The single-generation set-up, with lab-prepared mated adults, could not capture the impact of high temperatures prior to sexual maturation. A further possibility is seasonal climatic changes: our experimental conditions correspond to February temperatures in the field, when the average daily maximum temperature reaches 30.6°C at lowland. This is when the rainy season starts, driving an increase in *Drosophila* abundance and may serve as an annual reset of species’ distributions. The field study of O’Brien et al. occurred in April, which is 2°C lower in average temperature and average daily maximum. As the weather cools, mid- and low-elevation areas will become more suitable for the upland species to colonize. The relative importance of abiotic and biotic factors in affecting species distributions may change seasonally. Spatial and temporal data on species distributions are not yet available for our system but would provide a further test of our conclusions.

### Low variations in upper thermal limits

Upper thermal limits vary little among species (Hoffmann 2010) and have limited adaptive potential (van Heerwaarden and Sgrò 2021; Kellermann et al. 2012). Nevertheless, ecologically meaningful variations do exist, especially in tropical regions (Amundrud and Srivastava 2020; García-Robledo et al. 2016; von May et al. 2017; Pintanel et al. 2021), indicating ecological sorting or adaptation. Consistent with other studies (Goulet, Thompson, and Chapple 2017; Hangartner and Hoffmann 2016), we found that heat tolerance is a systematic trait manifested in critical temperatures, productivities at sub-sterile temperatures, recovered productivities, and locomotive responses. Modest variations in critical temperature may signify ecologically meaningful differences in overall performance under real and variable thermal conditions.

Such small thermal safety margins suggest a severe threat of biotic attrition in tropical lowlands (Colwell et al. 2008; Deutsch et al. 2008; Duarte et al. 2012; van Heerwaarden and Sgrò 2021). Laboratory-measured critical temperatures are sensitive to experimental conditions, making it difficult to relate the exact values to climatological means or maxima, and hence the threat of rising temperatures (Sinclair et al. 2016). In this context, our study benefits from a comparative approach, which reveals that upland species are already constrained by high temperatures at low elevations, and that a very small difference (about 1°C) distinguishes the upper thermal limits (*RTmax*s) of lowland and upland species. Given the low evolutionary potential of heat tolerance (Hoffmann, Chown, and Clusella-Trullas 2013), both lowland species and upland species are likely to be vulnerable to modest temperature increases across the elevation gradient. Thus, lowland biotic attrition and upland range contraction are likely to occur with future warming. This could lead to cascading effects in lowland communities and threaten endemic upland species on tropical mountains such as those in the Australian Wet Tropics (Freeman et al. 2018).

### Daily peak temperature as the main abiotic filter

Daily maxima can be more important than mean temperatures in structuring distributions (Lynch et al. 2014). In our study system, the mean temperature during the study season at our lowland sites is around 26°C. At this temperature, all studied species are close to their peak reproductive performance. However, upland and lowland sites differ greatly in the average daily maximum temperature and the daily duration that the temperature reaches or exceeds stressful levels for *Drosophila* reproduction (Supplementary Figure 7). In a preliminary experiment where populations were maintained for multiple generations in mixed-species cultures at constant 26°C, *D. pallidifrons* always out-numbered *D. pandora* (Jinlin Chen, data not shown), contrasting poor performance of *D. pallidifrons* at 24°C /28.5°C. Brief exposure to stressful thermal environments can have similar fitness costs to continuously stressful conditions (Saxon, O’Brien, and Bridle 2018). These results highlight the importance of considering daily temperature variations and extreme temperature events when studying species distributions and projecting responses to climate change (Kingsolver, Diamond, and Buckley 2013; Ma, Hoffmann, and Ma 2015).

### Limitations

Our laboratory environment has low variations in temperature, spatial structure, and diet, limiting the mechanisms organisms use to withstand thermal stress and competitive pressure. Behavioural thermoregulation allows species with distinctive fundamental thermal tolerance to exist in one area but using different microhabitats (Vives-Ingla et al. 2022). This spatial or temporal partitioning can also influence the competition between resource-sharing species (Bestelmeyer 2008; Porras et al. 2020). Furthermore, competition in reproduction and larval survival may be overrepresented in restricted space inside containers, while differences in abilities to forage, escape, and use alternative resources were not captured. Nevertheless, our thermal performance curves and monoculture experiments showed that lowland species could persist in abiotic conditions outside their observed range. It implies that biotic mechanisms, likely competitive exclusion, play a role in upland community composition. The match between reproduction limits of upland species and recorded air temperature in lowlands implies their lack of ability to mitigate the adverse impact of high temperature on natural populations, such as direct heat stress on larvae on rotten fruits or deficiency in foraging under heat.

Finally, between-population variation and local adaptation (Hoffmann, Anderson, and Hallas 2002), especially adaptation to abiotic environments at distribution boundaries (Peterson, Doak, and Morris 2019), should not be neglected if studying distributions at large spatial scales (e.g., across latitudes). The relatively restricted spatial scale of our study (within a mountain range), the use of mass-bred lines sourced from different elevations, and the limited plasticity and local adaptation documented for our study species (MacLean et al. 2019; O’Brien et al. 2017) mean that plastic and evolutionary response to thermal conditions is unlikely to complicate interpretation of our results.

### Conclusions

Tropical ecosystems host an exceptional diversity of endemic species (Laurance et al. 2011). Predicting their sensitivity to climate change is a high priority and requires an understanding of the proximate causes of current distributions as well as the interactions between thermal tolerance and other environmental factors, both biotic and abiotic. Our study contributes to the growing literature demonstrating that species are sensitive to high temperatures at the warm boundaries of their distributions in the tropics. In particular, we highlight the important role of daily maximum temperatures in structuring communities in tropical lowland sites, and the essential role of interspecific competition at upland sites. Our results imply the vulnerability of both upland and lowland communities to increasing temperatures. These results demonstrate that reliable predictions of the sensitivity of species to rising temperatures may not always be achieved using simple correlational approaches; instead, a detailed understanding of multiple thermal traits and interactions with other species may be needed.

## Supporting information

Supplementary results

## Acknowledgements

We thank Jan Hrček (Czech Academy of Sciences) and Megan Higgie (James Cook University, Townsville) and their research groups for their assistance establishing *Drosophila* laboratory cultures. Jan Hrček, Chia-Hua Lue, Nick Pardikes, Mélanie Thierry and the Oxford Fly group provided valuable advice and shared facilities. We thank Chris Terry, Mélanie Thierry, Benjamin Van Doren, Eleanor O’Brien and Mukhlish Jamal Musa Holle for advice on data analysis and comments on the manuscript. We thank three anonymous reviewers for their constructive comments. This work was supported by NERC grant NE/N010221/1 to OTL and a tuition grant from the China Scholarship Council to JC.

## Author Contributions

JC and OTL both contributed to the development of ideas. JC designed and conducted the experimental work. JC analysed the results and led the writing of the manuscript. OTL contributed to the writing. JC and OTL both contributed to the revision.

