## Supplementary results for "Climate and competition combine to set elevational distributions of tropical rainforest *Drosophila*"

**Supplementary Table 1.** Isofemale lines used to construct mass-bred lines of the rainforest *Drosophila* species.

| Species | Origin of cultured lines (yes/no) | | | | Lines for mass bred lines |
| --- | --- | --- | --- | --- | --- |
|  | K low^1^ | K high^1^ | P low^1^ | P high^1^ |  |
| *D. bunnanda* | yes | no | yes | no | KL87, KL134, KL127, PL114 |
| *D. pandora* | no | no | yes | no | PL17, PL21, PL01^2^ |
| *D. bipectinata* | yes | no | yes | no | KL84, KL43, PL85, PL20 |
| *D. pseudoananassae* | yes | yes | yes | no | KL19, KH25, PL30, KH42 |
| *D. sulfurigaster* | yes | yes | yes | yes | KL08, KH10, PL51, PH18 |
| *D. birchii* | yes | yes | yes | yes | KL22, KH26, PL122, PH169 |
| *D. pallidifrons* | no | yes | no | yes | KH20, KH69, PH183, PH184 |
| *D. simulans* | no | no | yes | no | PL45, PL34, PL42, PL43 |
| *D. pseudotakahashii* | no | yes | no | yes | PH171, KH71^3^ |

Note:

1. “K” is Kirrama, “P” is Paluma. “low” and “L” indicate low elevation sites. “high” and “H” indicate high elevation sites.
2. Only these three isofemale lines were available.
3. Two uncontaminated isofemale lines were used to construct the mass-bred population later than the other listed species before the competition experiments.

**Supplementary Table 2.** Distribution patterns quantified by *hIndex* and by the effect (coefficient) of elevation on the likelihood of detecting each species. The effects of mountain and the elevation X mountain interaction are included except for *D. bunnanda*, *D. pandora*, and *D. pseudotakahashii*, where it was not possible to include the interaction term and/or the mountain term in the statistical models because of their very skewed distributions (very few observations at two elevations in either or both mountain ranges).

| ***hIndex*** | **Species** | **Factor** | **Coefficient** | **P value** |
| --- | --- | --- | --- | --- |
| 0.00 | *D. bunnanda* | elevation | -64.0 | 0.99 |
| 0.07 | *D. pandora* | **elevation** | **-7.80** | **0.002** |
| 0.12 | *D. bipectinata* | **elevation** | **-3.05** | **0.02** |
|  |  | mountain | **1.83** | **0.001** |
|  |  | **elevation*mountain** | **-6.22** | **0.02** |
| 0.40 | *D. pseudoananassae* | elevation | -0.53 | 0.22 |
|  |  | mountain | 0.18 | 0.60 |
|  |  | **elevation*mountain** | **-4.82** | **<0.001** |
| 0.43 | *D. rubida* | elevation | -0.77 | 0.06 |
|  |  | mountain | -0.29 | 0.29 |
|  |  | elevation*mountain | 0.64 | 0.22 |
| 0.54 | *D. sulfurigaster* | elevation | -0.91 | 0.54 |
|  |  | mountain | 0.11 | 0.90 |
|  |  | elevation*mountain | -0.01 | 0.99 |
| 0.64 | *D. birchii* | elevation | 1.24 | 0.27 |
|  |  | mountain | 0.40 | 0.62 |
|  |  | elevation*mountain | -0.94 | 0.51 |
| 0.76 | *D. pallidifrons* | **elevation** | **2.90** | **<0.001** |
|  |  | mountain | 0.44 | 0.39 |
|  |  | elevation*mountain | -0.54 | 0.51 |
| 1.00 | *D. pseudotakahashii* | **elevation** | **12.5** | **<0.001** |

**Supplementary Table 3.** Pairwise comparison (Tukey test) of knockdown times at low temperature for the nine *Drosophila* species. BIP = *D. bipectinata*, BIR = *D.birchii*, BUN = *D. bunnanda*, PAL = *D. pallidifrons*, PAN = *D. pandora*, PSA = *D. pseudoananassae*, SIM = *D. simulans*, SUL = *D. sulfurigaster*. MEL = *D. melanogaster*.

|  | **Difference** | **ci_lower** | **ci_upper** | **Adjusted p value** |
| --- | --- | --- | --- | --- |
| BIR-BIP | -0.29 | -1.29 | 0.72 | 0.99 |
| BUN-BIP | -0.12 | -1.12 | 0.88 | 1.00 |
| **MEL-BIP** | **7.29** | **6.28** | **8.29** | **0.00** |
| PAL-BIP | -0.14 | -1.14 | 0.86 | 1.00 |
| PAN-BIP | 0.64 | -0.36 | 1.64 | 0.54 |
| PSA-BIP | -0.33 | -1.33 | 0.67 | 0.98 |
| **SIM-BIP** | **5.07** | **4.07** | **6.07** | **0.00** |
| SUL-BIP | -0.55 | -1.55 | 0.45 | 0.74 |
| BUN-BIR | 0.17 | -0.83 | 1.17 | 1.00 |
| **MEL-BIR** | **7.57** | **6.57** | **8.57** | **0.00** |
| PAL-BIR | 0.14 | -0.86 | 1.14 | 1.00 |
| PAN-BIR | 0.93 | -0.07 | 1.93 | 0.09 |
| PSA-BIR | -0.05 | -1.05 | 0.95 | 1.00 |
| **SIM-BIR** | **5.36** | **4.36** | **6.36** | **0.00** |
| SUL-BIR | -0.26 | -1.26 | 0.74 | 1.00 |
| **MEL-BUN** | **7.40** | **6.40** | **8.41** | **0.00** |
| PAL-BUN | -0.02 | -1.03 | 0.98 | 1.00 |
| PAN-BUN | 0.76 | -0.24 | 1.76 | 0.30 |
| PSA-BUN | -0.21 | -1.22 | 0.79 | 1.00 |
| **SIM-BUN** | **5.19** | **4.19** | **6.19** | **0.00** |
| SUL-BUN | -0.43 | -1.43 | 0.57 | 0.92 |
| **PAL-MEL** | **-7.43** | **-8.43** | **-6.43** | **0.00** |
| **PAN-MEL** | **-6.64** | **-7.64** | **-5.64** | **0.00** |
| **PSA-MEL** | **-7.62** | **-8.62** | **-6.62** | **0.00** |
| **SIM-MEL** | **-2.21** | **-3.22** | **-1.21** | **0.00** |
| **SUL-MEL** | **-7.83** | **-8.83** | **-6.83** | **0.00** |
| PAN-PAL | 0.79 | -0.22 | 1.79 | 0.26 |
| PSA-PAL | -0.19 | -1.19 | 0.81 | 1.00 |
| **SIM-PAL** | **5.21** | **4.21** | **6.22** | **0.00** |
| SUL-PAL | -0.40 | -1.41 | 0.60 | 0.94 |
| PSA-PAN | -0.98 | -1.98 | 0.03 | 0.06 |
| **SIM-PAN** | **4.43** | **3.43** | **5.43** | **0.00** |
| **SUL-PAN** | **-1.19** | **-2.19** | **-0.19** | **0.01** |
| **SIM-PSA** | **5.40** | **4.40** | **6.41** | **0.00** |
| SUL-PSA | -0.21 | -1.22 | 0.79 | 1.00 |
| **SUL-SIM** | **-5.62** | **-6.62** | **-4.62** | **0.00** |

**Supplementary Figure 1.** Measuring reproductive thermal performance. A) Procedures and schedule of measurements. B) Changes in productivities from 1^st^ – 2^nd^ day to 7^th^ – 8^th^ day at different temperatures. BIP = *D. bipectinata*, BIR = *D.birchii*, BUN = *D. bunnanda*, PAL = *D. pallidifrons*, PAN = *D. pandora*, PSA = *D. pseudoananassae*, SIM = *D. simulans*, SUL = *D. sulfurigaster*.

A)

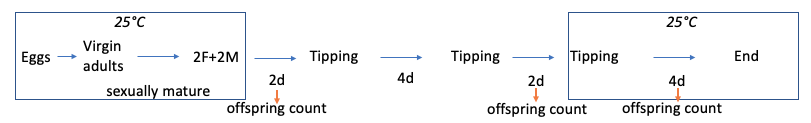

B)

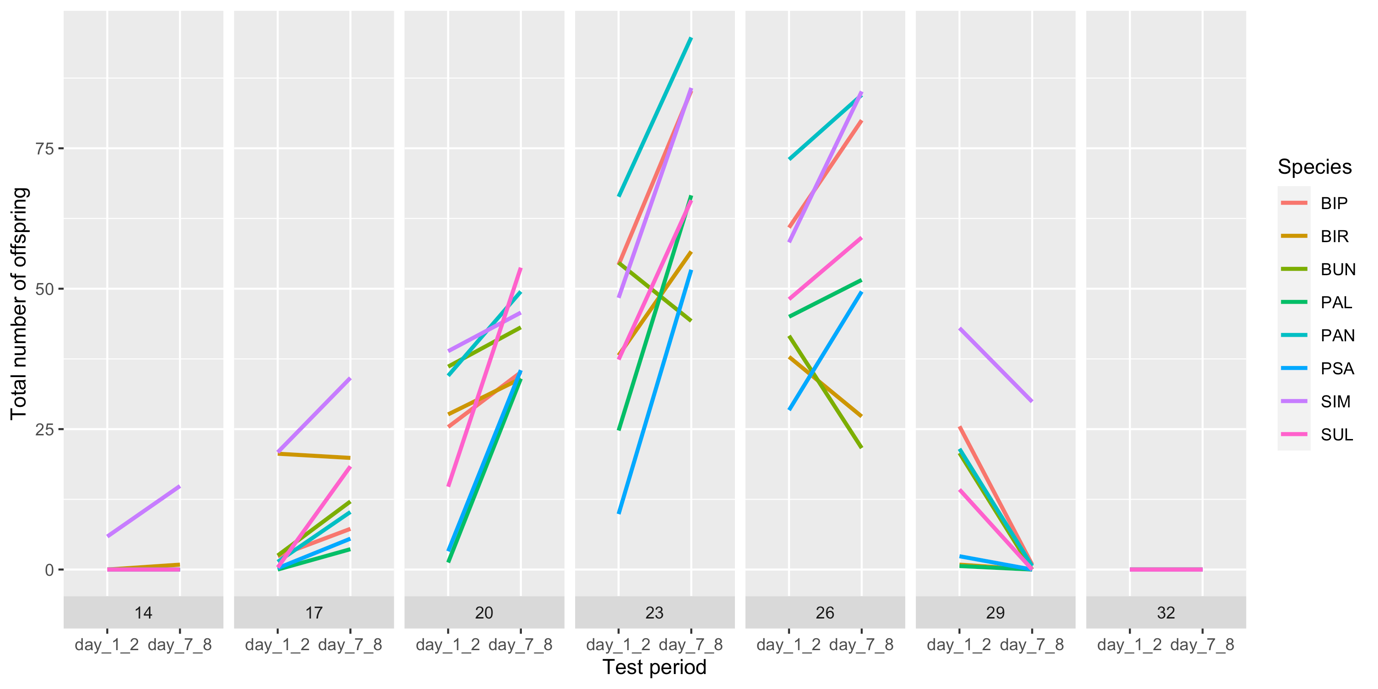

**Supplementary Figures 2.** Diagnostics of fitted thermal performance curves. A) Comparison of observed and fitted values of productivity. B) Residuals and standardized residuals plot. C) Observed daily productivity (points), fitted thermal performance curve (blue line) and its 95% credible interval for individual *Drosophila* species. To calculate the 95% credible interval, 200 sets of parameters were randomly drawn from their posterior distributions, the estimated values were calculated based on the parameters, and we extracted the upper and lower bounds of the 95% quantile of the estimated values.

A)

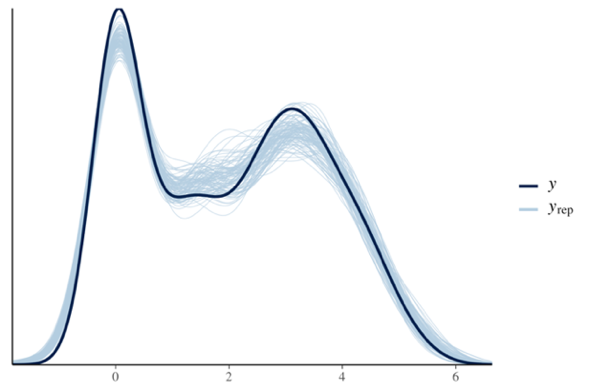

B)

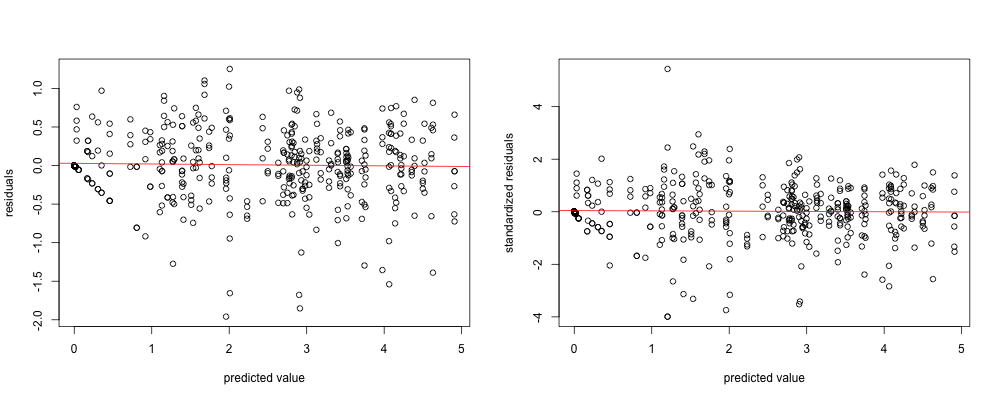

C)

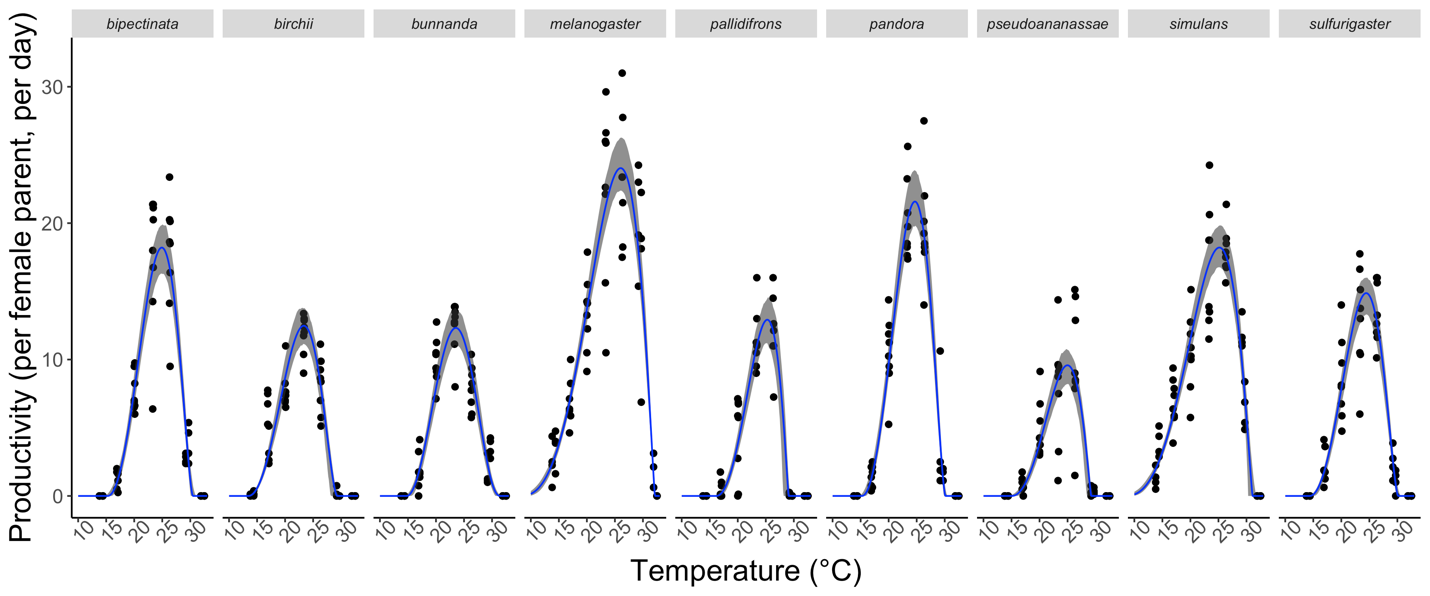

**Supplementary Figures 3.** A) The phylogenetic tree of all *Drosophila* species mentioned in this study. B) The ultrametric phylogenetic tree of the seven species used in the regression analysis of thermal traits. C) The process to create the ultrametric tree from the DrosoPhyla dataset (Finet et al., 2021).

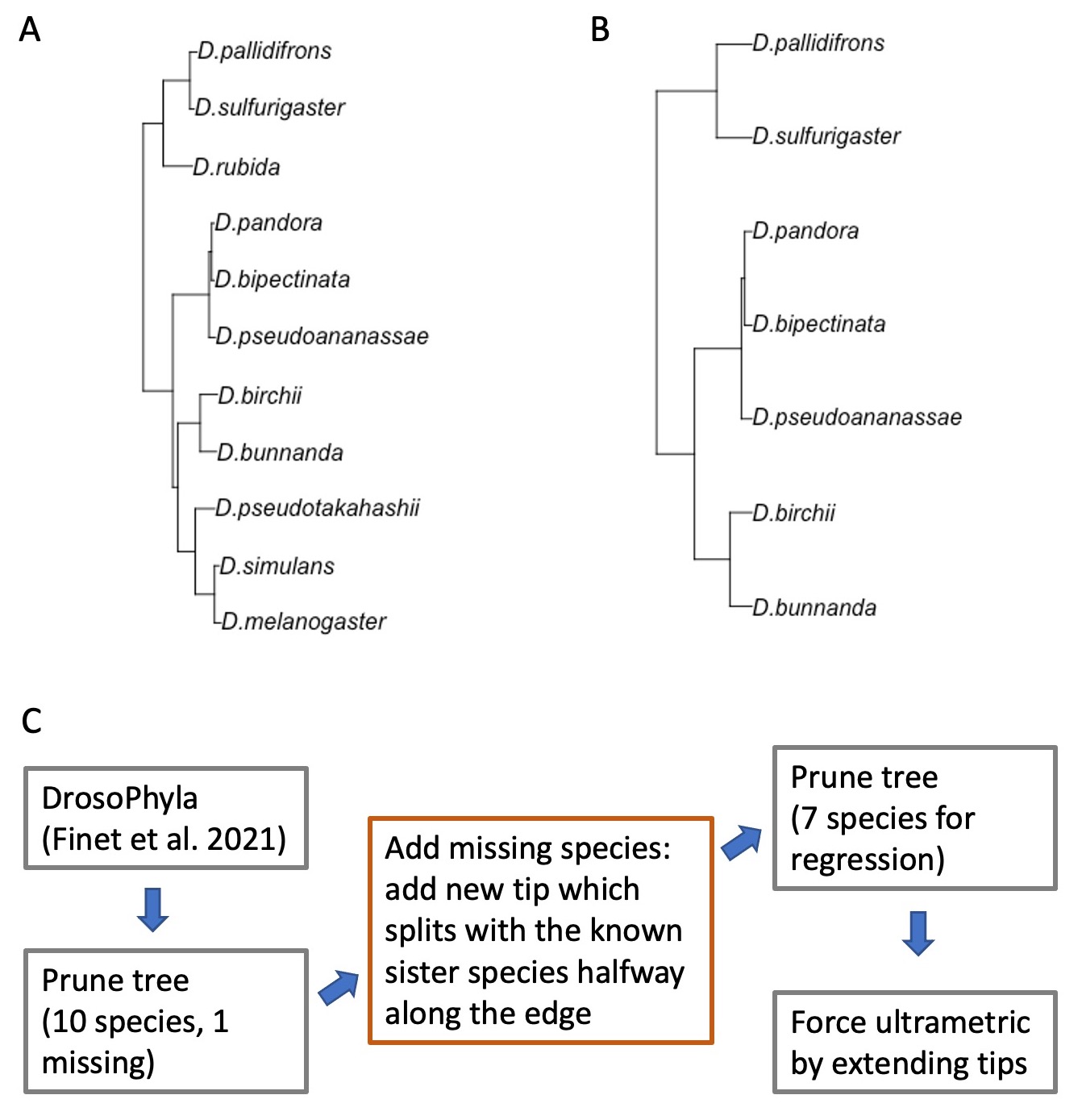

**Supplementary Figure 4.** Field and experimental temperatures of lowland (A) and upland (B) sites at Kirrama and Paluma in February 2017. Each grey line shows the hourly temperature in one day. The blue lines show the average temperature recorded at the time of the day. Temperature regimes used in short-term and long-term competition experiments are indicated by red lines, mimicking the day/night fluctuation of temperatures in the field.

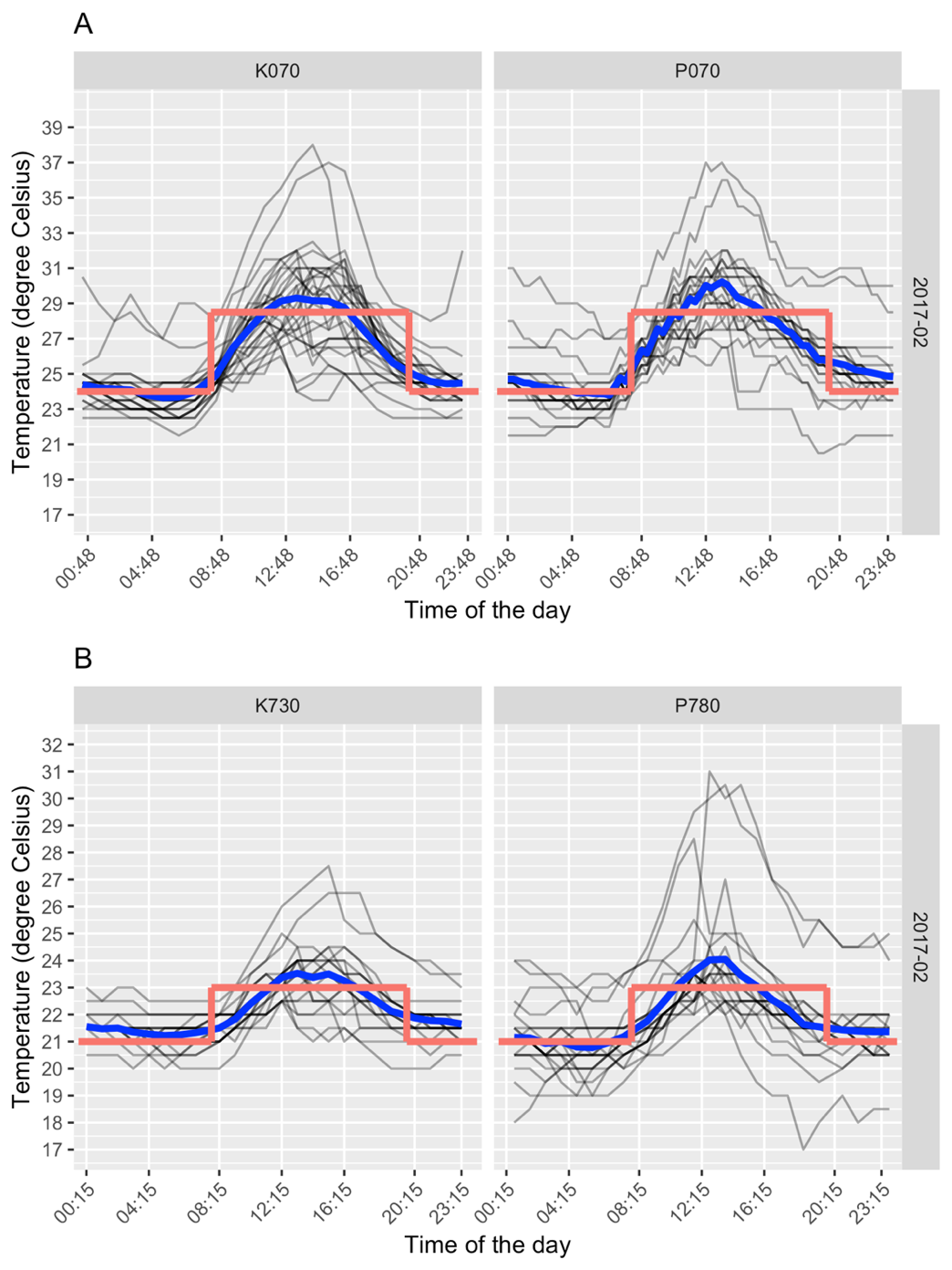

**Supplementary Figure 5.** Diagnostics of the fitted model for short-term competition (Beverton-Holt model) in the cool, upland (A) and warm, lowland (B) treatments, comparing the observed offspring counts and the fitted values. (C) Comparison of the observed population size and the fitted values from the regression for the long-term competition experiment.

A)

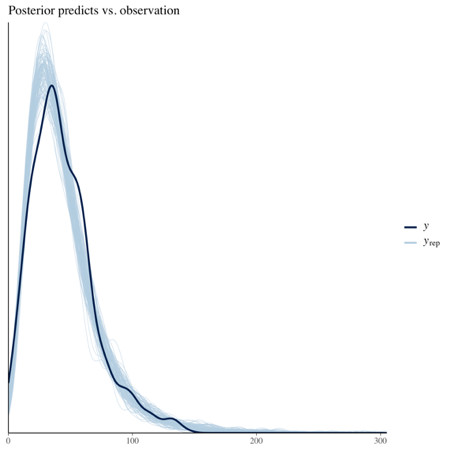

B)

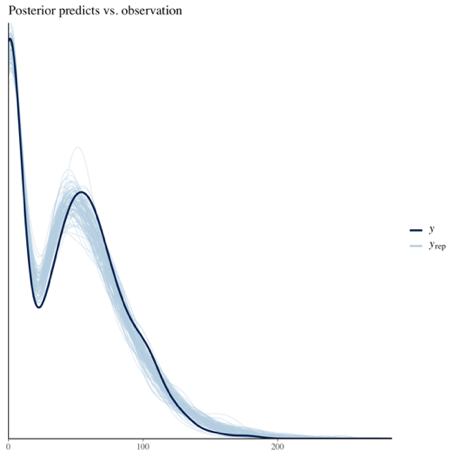

C)

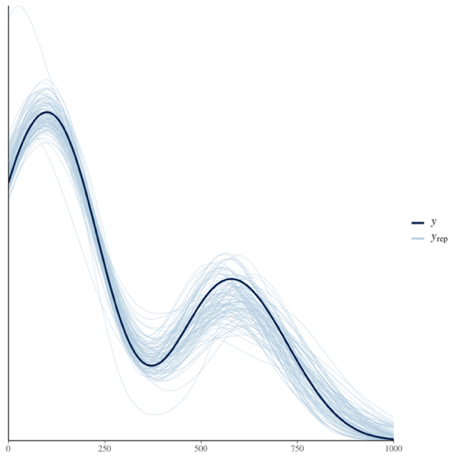
**Supplementary Figure 6.** Observed two-day productivity (points) from 4 pairs of parents and their fitted values (lines) in the short-term pairwise competition experiment. Upland species: PST = *D. pseudotakahashii*, PAL = *D. pallidifrons*. Elevation generalist: SUL = *D. sulfurigaster*. Lowland species: BIP = *D. bipectinata*, PAN = *D. pandora*.

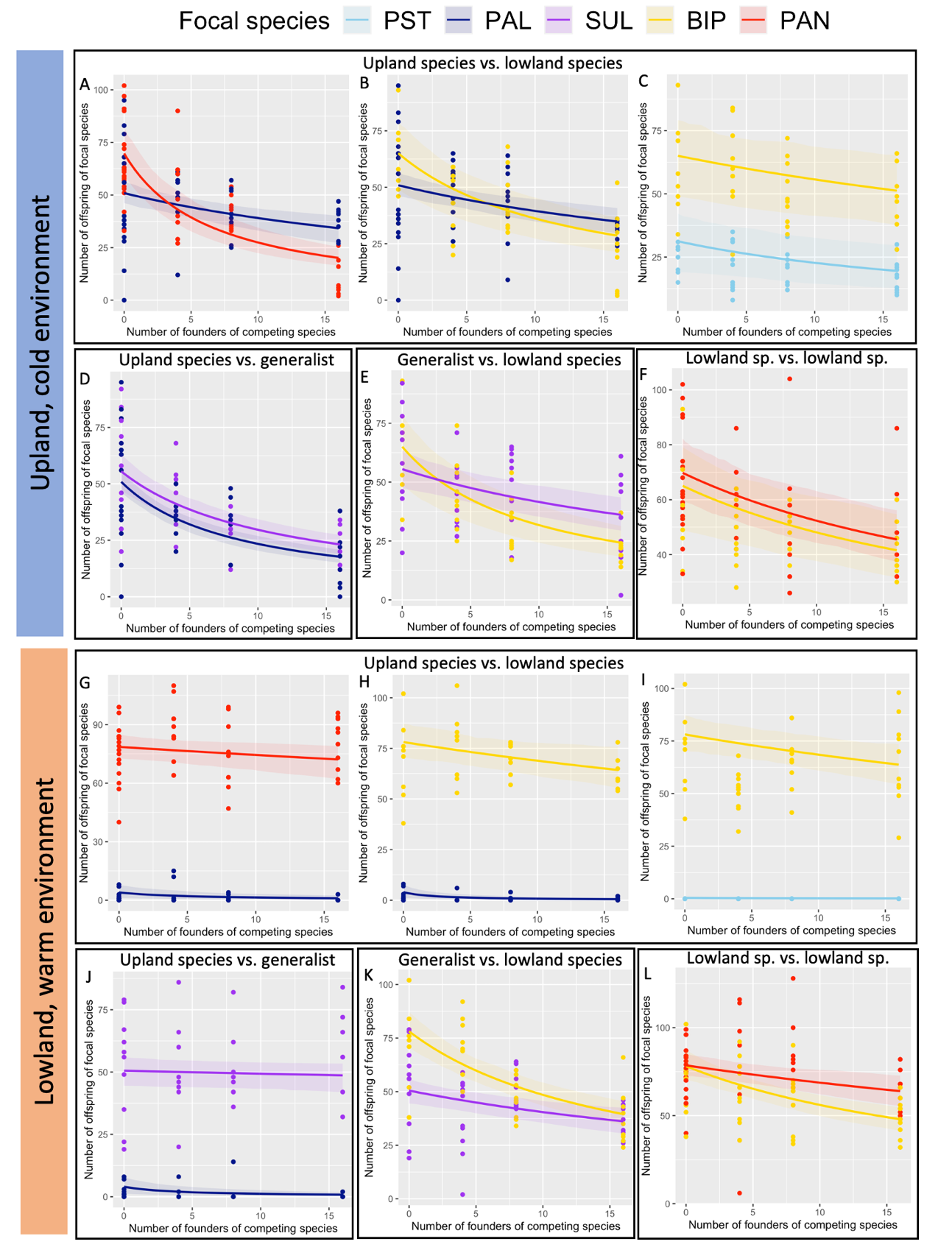

**Supplementary Figure 7. (**A) Average of daily maximum temperature in each month. The horizontal dark red line is 29°C, a known stressful temperature for reproduction. (B) Average hours per day that the temperature exceeds 29°C. Blue, green and red colours indicate sites at high, medium and low elevations, respectively. Solid and dashed lines represent sites at Kirrama and Paluma, respectively.

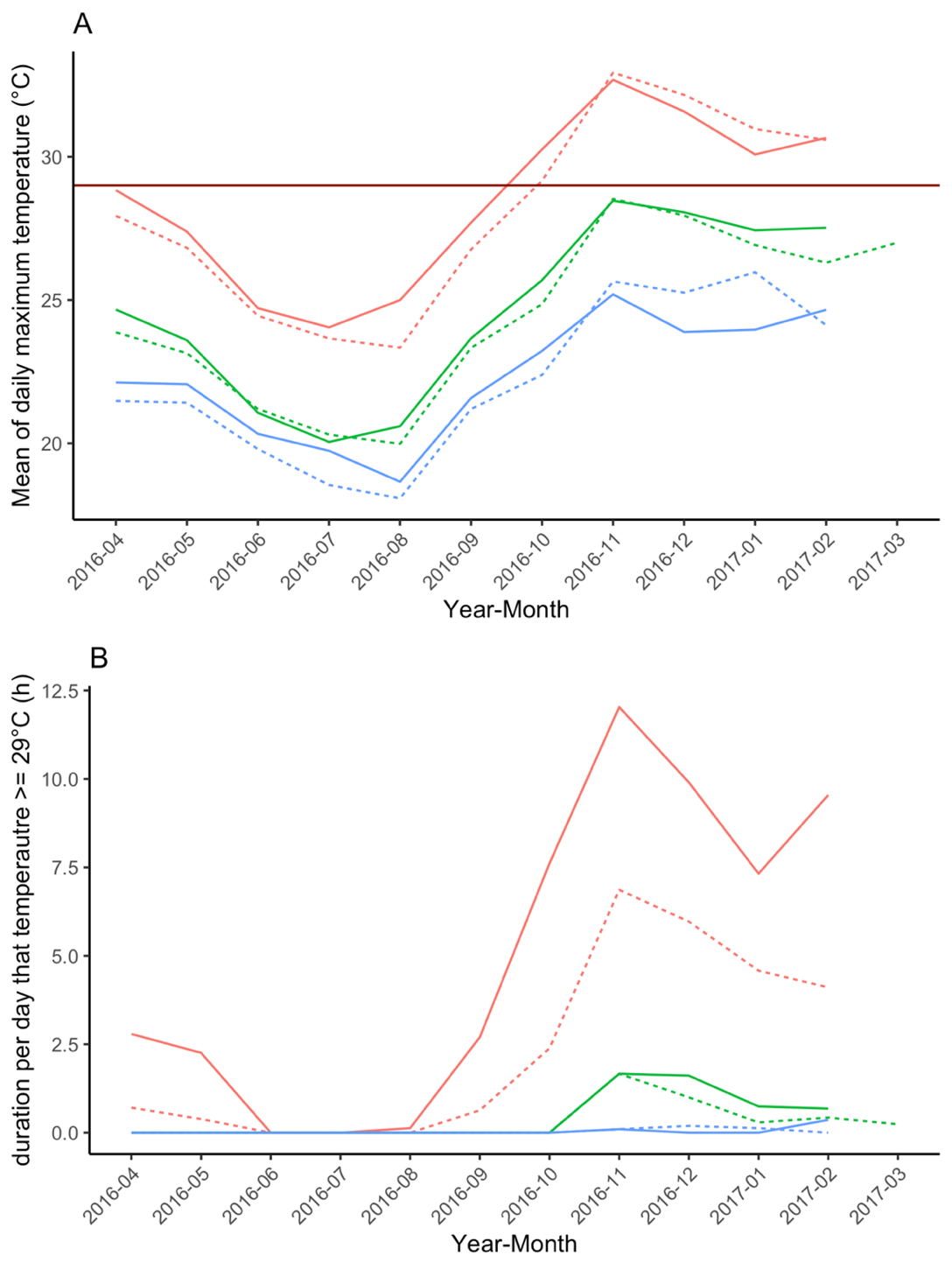
